# Exploration of molecular mechanism underlying the self-flocculation of *Zymomonas mobilis* through comparative omics analyses and experimental validations for developing robust production systems

**DOI:** 10.1101/2021.06.18.448934

**Authors:** Lian-Ying Cao, Yongfu Yang, Chen-Guang Liu, Yunhao Chen, Xue Zhang, Xia Wang, Juan Xia, Shihui Yang, Feng-Wu Bai

**Affiliations:** State Key Laboratory of Microbial Metabolism, Joint International Research Laboratory of Metabolic & Developmental Science of the Ministry of Education, and School of Life Science and Biotechnology, Shanghai Jiao Tong University, Shanghai 200240, China; State Key Laboratory of Biocatalysis and Enzyme Engineering, Environmental Microbial Technology Center of Hubei Province, and School of Life Science, Hubei University, Wuhan 430062, China

**Keywords:** *Zymomonas mobilis*, Self-flocculation of bacterial cells, Comparative genomics and transcriptomics, Molecular docking simulations, Biosynthesis of cellulose microfibrils

## Abstract

*Zymomonas mobilis* metabolizes sugar through the Entner-Doudoroff pathway with less ATP generated for lower biomass accumulation and more substrate to product formation with improved yield, since ATP is dissipated predominately through growth for intracellular energy homeostasis, making it a platform to be engineered as microbial cell factories, particularly for producing bulk commodities with major cost from feedstock consumption. ZM401, a self-flocculating mutant, presents advantages for production including cost-effective biomass recovery through gravity sedimentation, self-immobilization within bioreactors for high cell density to improve productivity and enhanced tolerance to environmental stresses for high product titers, but molecular mechanism underlying this phenotype is largely unknown. In this work, we sequenced and assembled the genome of ZM401 to explore genetic basis for the self-flocculation of the bacterial cells through comparative genomic and transcriptomic analyses, molecular docking simulations for enzymes encoded by functional genes and their substrates/activators, and experimental validations. Our results demonstrated that the single nucleotide deletion in ZMO1082 disrupted its stop codon for the putative gene being fused with ZMO1083, which created an exciting gene encoding the subunit A of the bacterial cellulose synthase with unique function for synthesizing cellulose microfibrils to flocculate the bacterial cells, and the single nucleotide mutation in ZMO1055 compromised the function of bifunctional diguanylate cyclase/phosphodiesterase encoded by the gene on the degradation of c-di-GMP for its intracellular accumulation to activate the cellulose biosynthesis. These discoveries are significant not only for optimizing the self-flocculation of *Z. mobilis*, but also engineering other bacteria with the self-flocculating phenotype for robust production.

## Introduction

Compared to the Embden-Meyerhof-Parnas (EMP) pathway commonly employed by other species, the ethanologenic bacterium *Zymomonas mobilis* metabolizes sugar through the Entner-Doudoroff (ED) pathway with much less ATP generated (Kalnenieks 2006), which consequently reduces its biomass accumulation, since ATP is dissipated predominately through cell growth for intracellular energy homeostasis as observed in ethanol fermentation with the brewing yeast *Saccharomyces cerevisiae* (de Kok et al. 2012). As a result, more sugar can be directed to product formation with improved yield, which is very significant for producing bulk commodities such as ethanol as a biofuel with major cost from feedstock consumption (Gombert and van. Maris 2015). On the other hand, the bacterial cells are much smaller than the yeast cells, and thus a high specific surface area is available for sugar uptake, which, together with the unique ED pathway, forms a catabolic highway for carbon metabolism (Rutkis et al. 2016; Sprenger 1996).

Moreover, *Z. mobilis* can be engineered to metabolize pentose sugars released from the hydrolysis of hemicelluloses in lignocellulosic biomass through the isomerase pathway without the challenge of cofactor imbalance, an intrinsic disadvantage in engineering *S. cerevisiae* for the same purpose (Zhang et al. 1995; Gopinarayanan and Nair 2019). These merits make the bacterium a platform to be engineered for biorefinery to produce biofuels and bio-based chemicals (He et al. 2014).

ZM401, a self-flocculating mutant developed through chemical mutagenesis by nitrosoguanidine treatment from ZM4, the unicellular model strain of *Z. mobilis*, presents many advantages for production. The self-flocculated bacterial cells can settle down quickly through gravity sedimentation from fermentation broth for biomass recovery at low cost (Movie 1) without a necessity for centrifugation, a common practice for harvesting unicellular cells in industry with high capital investment on centrifuges and intensive energy consumption on their running. Unlike packed-bed and fluidized-bed reactors for chemical reactions, bioreactors for microbial conversion are characterized by low cell density, particularly under continuous culture due to an auto-balance between the growth of microbial cells and their washing-out with the effluent for chemostat conditions (Novick and Szilard 1950). Cell immobilization within bioreactors can address this challenge, but supporting materials are needed for immobilizing unicellular cells, which is not acceptable for producing bulk commodities with marginal revenues because of additional cost associated with the consumption of supporting materials. When microbial cells self-flocculate, they can be immobilized within bioreactors without consumption of supporting materials, which has been highlighted in continuous ethanol fermentation using the self-flocculating yeast (Zhao and Bai 2009).

The most important trait for microbial cells to perform robust production is their tolerance to stresses (Gong et al. 2017). Product inhibition is one of them, since high product titer has been pursued endlessly to save energy consumption in downstream product recovery, and in the meantime to reduce wastewater discharge. Toxicity from by-products is another, which is one of the biggest challenges for biorefinery of lignocellulosic biomass, since many toxic by-products are released during its pretreatment (Ling et al. 2014). Although specific stress response has been intensively studied, general stress response (GSR) is more preferred, because multiple stresses are always developed simultaneously under industrial production conditions (Guan et al. 2017).

Quorum sensing (QS) is a major mechanism underlying GSR, particularly in bacterial cells through secreting signal molecules as public goods to collectively fight against stressful environments for survival, but high cell density is a prerequisite for signal molecules to approach their thresholds to trigger QS (Mukherjee and Bassler 2019). We have observed that the self-flocculating ZM401 is more tolerant to major inhibitors in the hydrolysate of lignocellulosic biomass including furfural, hydroxymethylfurfural (HMF), acetic acid and vanillin (Zhao et al. 2014), and speculated the underlying mechanism would be enhanced QS, since the self-flocculation of ZM401 characterized by cell-to-cell contact is the upper limit for localized high cell density, and QS could be triggered more effectively.

However, molecular mechanism underlying the self-flocculation of *Z. mobilis* remains largely unknown. Recently, chemial basis for the self-flocculation of ZM401 was identified to be cellulose microfibrils (Xia et al. 2018), and the genome of ZM4 was sequenced previously and updated twice (Seo et al. 2005; Yang et al. 2009a; Yang et al. 2018), providing references for comparative studies on the genome of ZM401 to explore molecular mechanism underlying its self-flocculating phenotype, which is significant not only for optimizing the self-flocculation of *Z. mobilis* at molecular levels through rational design, but also for engineering other bacteria with the phenotype for robust production.

## Results

### Genetic basis for the self-flocculation of ZM401

Combining the data collected from the long-read and short-read sequencing, we assembled genome for ZM401 to explore the genetic basis for the self-flocculation of the bacterial cells through comparative genomics analysis between ZM401 and ZM4. The genome contains one circular chromosome of 2,058,754 bp and four circular plasmids of 32,791 bp (401_pZM32), 33,006 bp (401_pZM33), 36,494 bp (401_pZM36) and 39,266 bp (401_pZM39), respectively. With the ZM4 genome information from the NCBI database as the reference (Yang et al. 2018), we annotated the genome of ZM401.

One nucleotide deletion was detected in the circular chromosome of ZM401, but the sizes of its four circular plasmids are same as those in ZM4. We identified totally 35 single nucleotide variations (SNVs), but no significant structure change was observed in the genome of ZM401. For the 35 SNVs, one nucleotide deletion occurred in the putative gene ZMO1082, and 34 single nucleotide polymorphisms (SNPs) occurred in other genes and intergenic regions as well (Table S1), which were verified by Sanger sequencing. However, the Sanger sequencing verification confirmed that the SNP within ZMOp32×018 in the plasmid pZM32 at the site of 1789 and another SNP within the intergenic region in the circular chromosome at the sites 1614973 for ZM401 are exactly same as the nucleotides at the site of 1789 and 1614974 for ZM4, whose genome is one nucleotide larger than that of ZM401. Therefore, the genome sequence at the database for ZM4 might be wrong at the two sites. As a result, we confirmed totally 33 SNVs in the genome of ZM401, of which 31 are within genes, and 2 are within untranslated regions (UTRs).

For the 31 SNVs in genes with confirmed or putative functions, one is nucleotide deletion in ZMO1082, a putative gene, and all others are SNPs. For the 30 SNPs, 17 of them are non-synonymous mutations (Table 1), and others are synonymous mutations. Two SNPs within the UTRs are located at the upstream of ZMO0914 and ZMO1909, respectively.

**Table 1.**
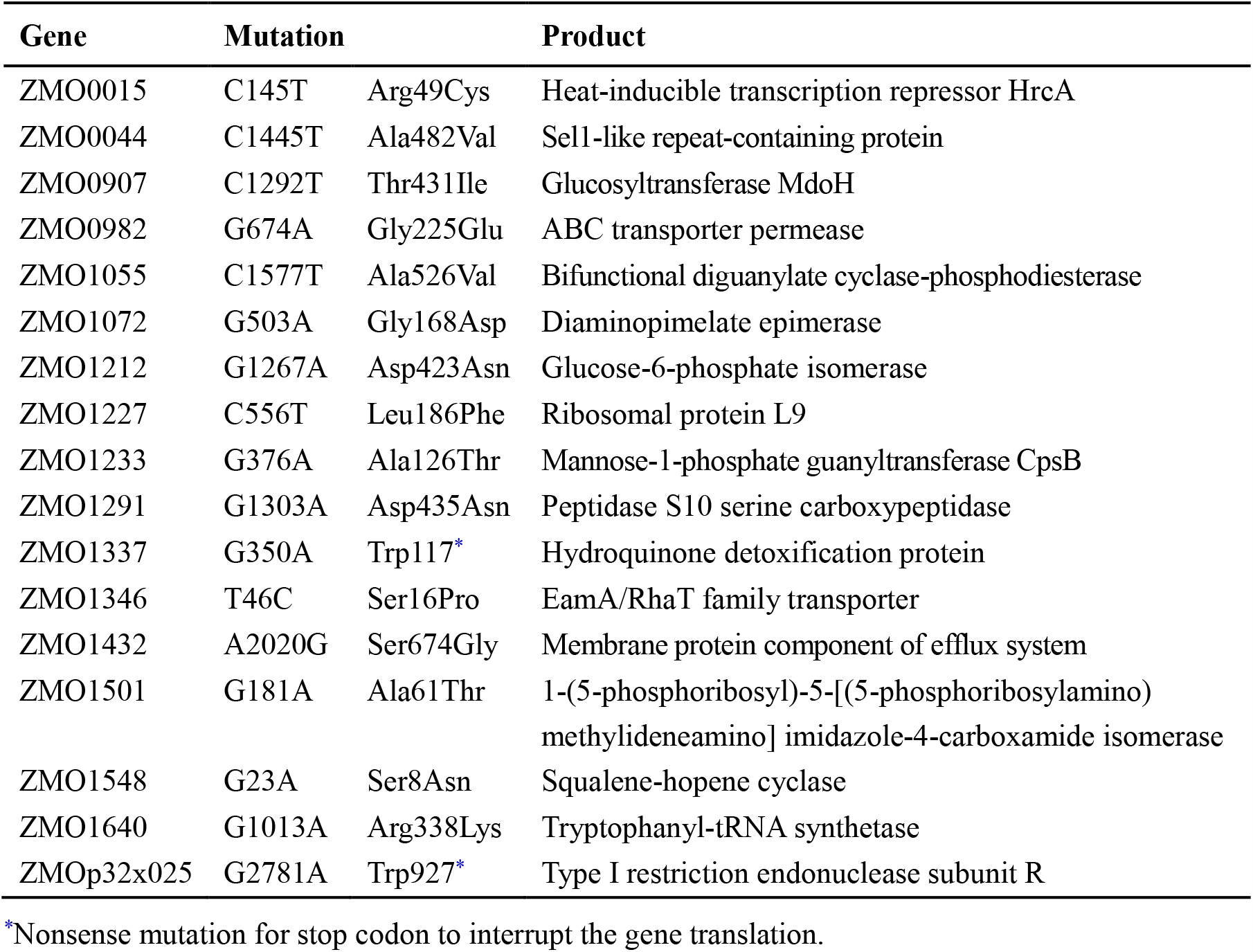
Non-synonymous mutations for genes in the genome of ZM401.

### Comparative transcriptomic analysis between ZM401 and ZM4

The RNA-seq analysis demonstrated that totally 74 genes were differentially expressed in ZM401 (63 up-regulated and 11 down-regulated), compared to those in ZM4 under same culture conditions (Fig. S1 and Table S2). For the 18 genes with mutations at protein level in ZM401, only ZMO1082 with the nucleotide deletion was significantly up-regulated. We therefore performed gene ontology enrichment for those differentially expressed genes to explore their functions (Fig. S2).

For those up-regulated genes, the largest set is for cell components, in particular cell membranes, followed by the set for biological processes associated with major cell wall components such as the synthesis of β-glucans including cellulose and the activation of glucose to form uridine 5′-diphosphoglucose (UDP-glucose) as substrate for cellulose biosynthesis, and the set for molecular functions such as transmembrane activities, which support our hypothesis for the self-flocculation of ZM401: the direct interactions of extracellular polysubstance(s) synthesized predominantly under the catalysis of membrane-embedded enzymes such as cellulose synthase for cellulose biosynthesis.

On the other hand, functions of the 11 down-regulated genes are predominately related to development and metabolism including DNA biosynthesis and replication as well as oxidoreductase activities, which are significantly different from functions observed for the up-regulated genes, indicating their less impact on the self-flocculation of ZM401. Theoretically, the self-flocculation of microbial cells might compromise their growth and metabolism to some extent due to a potential impact of the morphology on nutrient uptake from medium, although this phenomenon seems not significant for ZM401.

### Impact of mutant genes on the transcription of other genes

Using the software DOOR^2^ and STRING (Mao et al. 2009; Szklarczyk et al. 2019), we searched operons for those significantly up-regulated genes, and predicted protein-to-protein interactions. As a result, ZMO1082, ZMO1083, ZMO1084 and ZMO1085 were suggested to be an operon (Figs. S3 and S4).

Although many genes in the genome of *Z. mobili*s are still putative, previous bioinformatics analysis indicated that ZM4 contains a bacterial cellulose synthase operon (*bcs*) comprised of ZMO1083, ZMO1084 and ZMO1085, encoding membrane-embedded cellulose synthase composed of subunits A, B and C (BcsA, BcsB and BcsC), respectively (Jeon et al. 2012). BcsA is the catalytic subunit for cellulose biosynthesis, and BcsB is a large periplasmic protein that is anchored to inner membrane guiding synthesized cellulose exported across the periplasm toward outer membrane, which are frequently fused together (Omadjela et al. 2013), and missense mutation in BcsA can change cellulose biosynthesis (Salgado et al. 2020).

ZMO1082 is a putative gene in ZM4, encoding a peptide with 67 amino acids only, which is too small and less likely functional, but its mutation in ZM401 created by the deletion of one thymine from the nine repetitive thymines coincidently affected ZMO1083, since such a mutation resulted in a frameshift event, and consequently disrupted the stop codon of ZMO1082 (TGA), making it fused with ZMO1083 to create a new gene ZMO1083/2 (**Fig. 1a**). With 60 more amino acids incorporated, the catalytic function of the BcsA was altered significantly for more efficient biosynthesis of cellulose, which was further developed as cellulose microfibrils to flocculate the bacterial cells **(Fig. 1b)**.

**Fig. 1.**
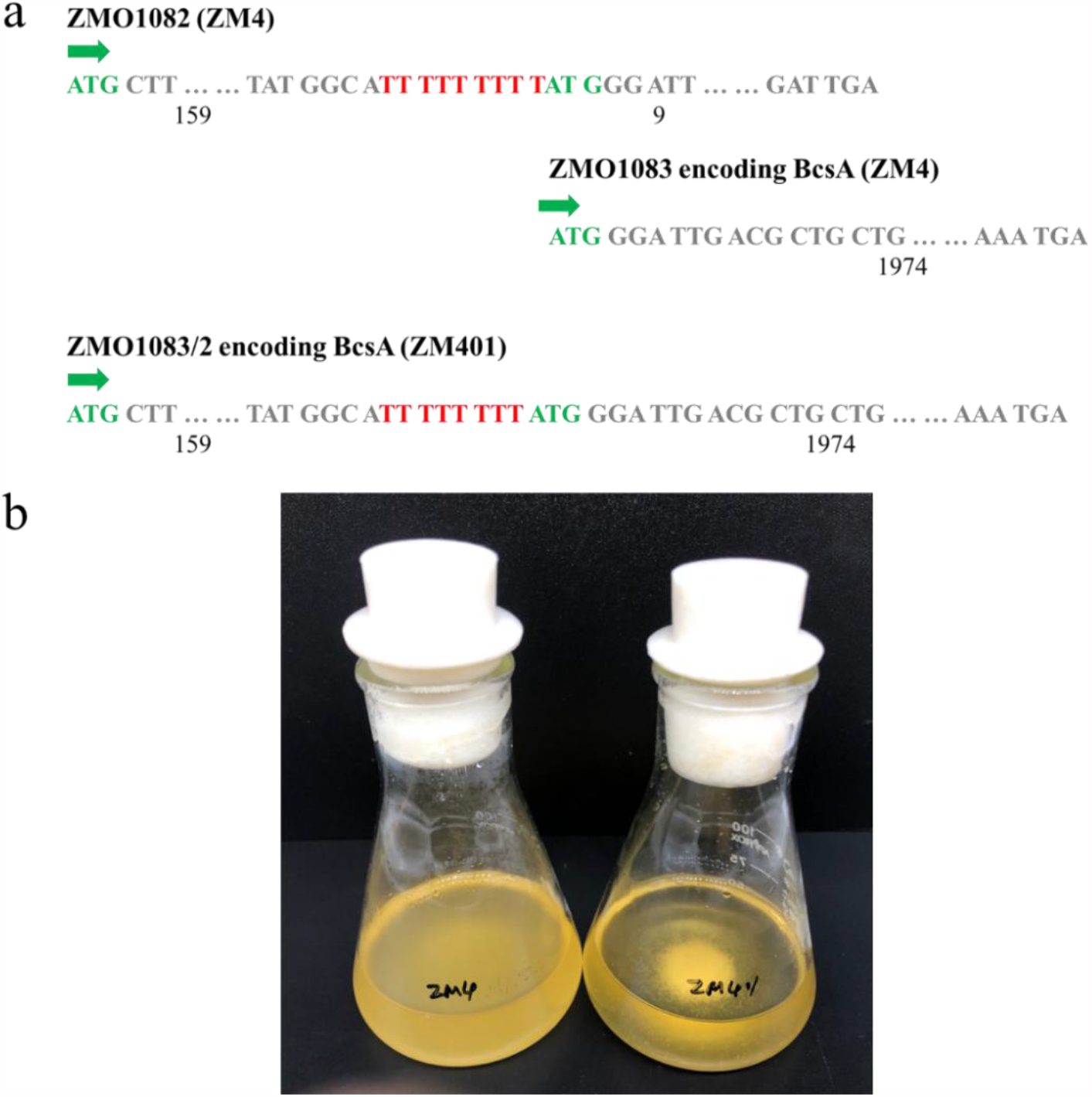
Nucleotide sequences for the putative gene ZMO1082 and its neighboring gene ZMO1083 in ZM4 and their fusion for the new gene ZMO1083/2 in ZM401 (a), and comparison for the morphologies of the non-flocculating ZM4 and self-flocculating ZM401 (b). The dynamic gravity sedimentation of ZM401 is highlighted in Movie 1.

Cyclic diguanylic acid (c-di-GMP) is a second messenger for signal transduction in bacteria to regulate intracellular processes including cellulose biosynthesis (Jenal et al. 2012; Ross et al. 1987). Bacteria synthesize c-di-GMP from guanosine triphosphate (GTP) under the catalysis of diguanylate cyclase, but phosphodiesterase can degrade c-di-GMP (Nesbitt et al. 2015). ZMO1055 in ZM4 is predicted to encode a bifunctional diguanylate cyclase/phosphodiesterase with the conserved motifs GGDQF and EAL, catalyzing the synthesis and degradation of c-di-GMP, respectively. Although the transcription level of ZMO1055 didn’t change significantly in ZM401, the amino acid mutation created by the SNP is located at the EAL domain, which could compromise its catalytic activity on c-di-GMP degradation for more accumulation of the messenger molecule to regulate intracellular processes effectively, particularly cellulose biosynthesis for the self-flocculation of the bacterial cells. The up-regulation of the whole *bcs* operon in ZM401 partly supported this speculation.

### Molecular docking for key enzymes and their substrates/activators

Based on the protein structures deposited at the Worldwide Protein Data Bank for the BcsA-BcsB complex in *Rhodobacter sphaeroides* (Morgan et al. 2013), we performed structural modelling using the software SWISS-MODEL (Waterhouse et al. 2018) for proteins encoded by ZMO1083/2 in ZM401 and ZMO1083 in ZM4, and further simulated the docking of substrate UDP-glucose and c-di-GMP with the enzymes, respectively.

**Fig. 2** indicated that active sites on protein encoded by ZMO1083 in ZM4 for binding with UDP-glucose and c-di-GMP are too close for them to bind simultaneously. However, protein encoded by ZMO1083/2 in ZM401 has two additional transmembrane regions at its N-terminus, which distance the site for UDP-glucose binding from the central cavity to the bottom of the beta-barrel, eliminating the overlap binding for c-di-GMP. Moreover, the site for c-di-GMP binding on the protein encoded by ZMO1083/2 in ZM401 is closer to the gate-loop than that encoded by ZMO1083 in ZM4 for more effective binding of the messenger molecule. These structural modifications not only enhance the biosynthesis of cellulose in ZM401, but also the development of cellulose microfibrils to flocculate the bacterial cells.

**Fig. 2.**
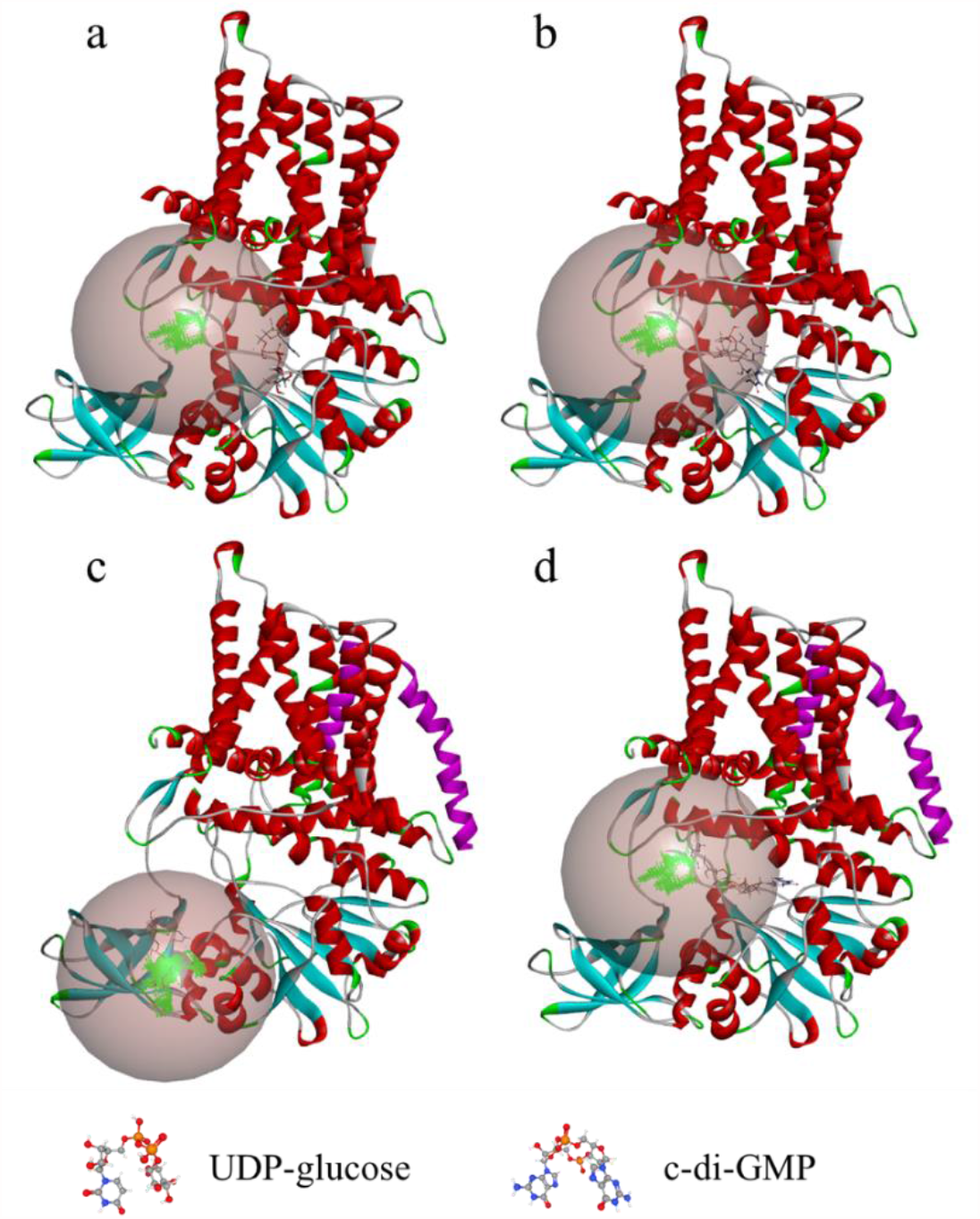
Molecular docking simulations for UDP-glucose and c-di-GMP with BcsA encoded by ZMO1083 in ZM4 (a and b) and ZMO1083/2 in ZM401 (c and d). Bright purple: mutation in the EAL domain, cinnamic ball: region with low energy for recruiting UDP-glucose and c-di-GMP, and green pocket: active sites.

The EAL domain is conserved in Gram-negative bacteria including *Z. mobilis* for catalyzing c-di-GMP degradation (Jenal et al. 2012). The mutation of Ala526Val on ZMO1055 in ZM401 is located at the EAL domain of the bifunctional diguanylate cyclase/phosphodiesterase encoded by the gene, which could affect its binding with c-di-GMP. To test this hypothesis, we performed structure modelling for proteins encoded by ZMO1055 in ZM401 and ZM4, respectively **(Fig. 3)**.

**Fig. 3.**
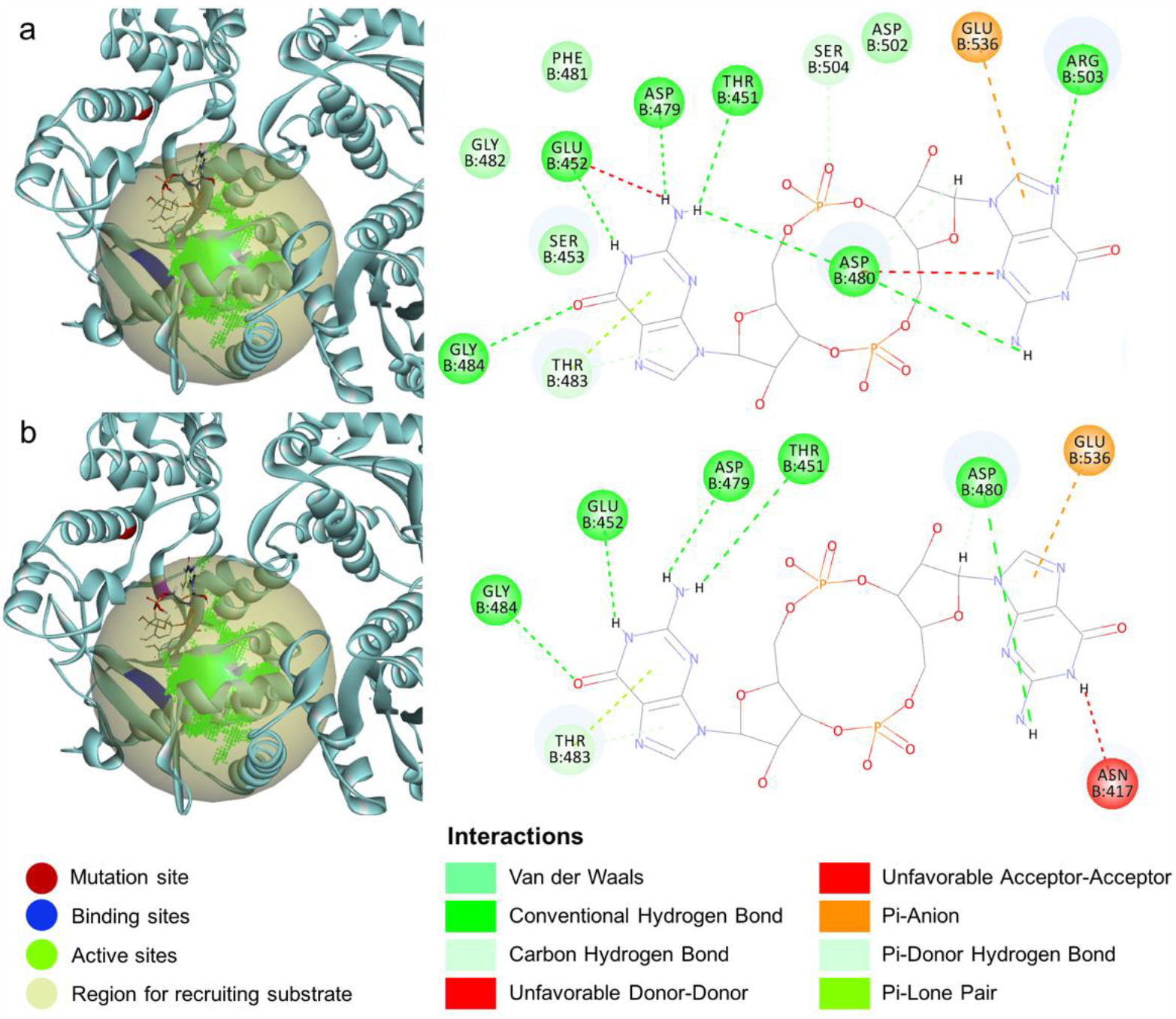
Molecular docking simulations for c-di-GMP with the active center of the EAL in the bifunctional enzyme (diguanylate cyclase/phosphodiesterase) encoded by ZMO1055 in ZM4 (a) and ZM401 (b), respectively.

The studies indicate that the single amino acid mutation cannot change the protein structure substantially, but molecular docking simulations show that the putative pocket for accommodating c-di-GMP is close to the mutation site, which might affect the binding of c-di-GMP to the EAL domain, and consequently compromise its degradation. On the other hand, the mutation Ala526Val might affect linkage between ASP480 and c-di-GMP through hydrogen bonds. In ZMO1055 from ZM4, ASP480 cyclizes with c-di-GMP, stabilizing the docking to make the EAL domain more effective for its degradation, but the docking of c-di-GMP into the EAL domain in ZM401 is less stable, since ASP480 links with c-di-GMP through one hydrogen bond only. As a result, the degradation of c-di-GMP could be compromised in ZM401 for its intracellular accumulation to enhance the synthesis of cellulose microfibrils to flocculate the bacterial cells (Morgan and McNamara 2014).

### Experimental validations

When either ZMO1082 or ZMO1083 was deleted from ZM401, the chord length detected by the focused beam reflectance measurement (FBRM) system decreased from 425.6 μm to the undetectable level for unicellular bacterial cells, indicating the self-flocculating phenotype of ZM401 was disrupted completely, but when ZMO1083/2 was overexpressed in ZM4, no self-flocculation was observed on the mutant (**Fig. 4a**), indicating that ZMO1083/2 is a prerequisite, and both ZMO1082 with the nucleotide deletion and ZMO1083 without any mutation are needed for the gene to be functional on synthesizing cellulose microfibrils to flocculate the bacterial cells, but ZMO1083/2 alone is not sufficient for ZM4 to develop the self-flocculating phenotype.

**Fig. 4.**
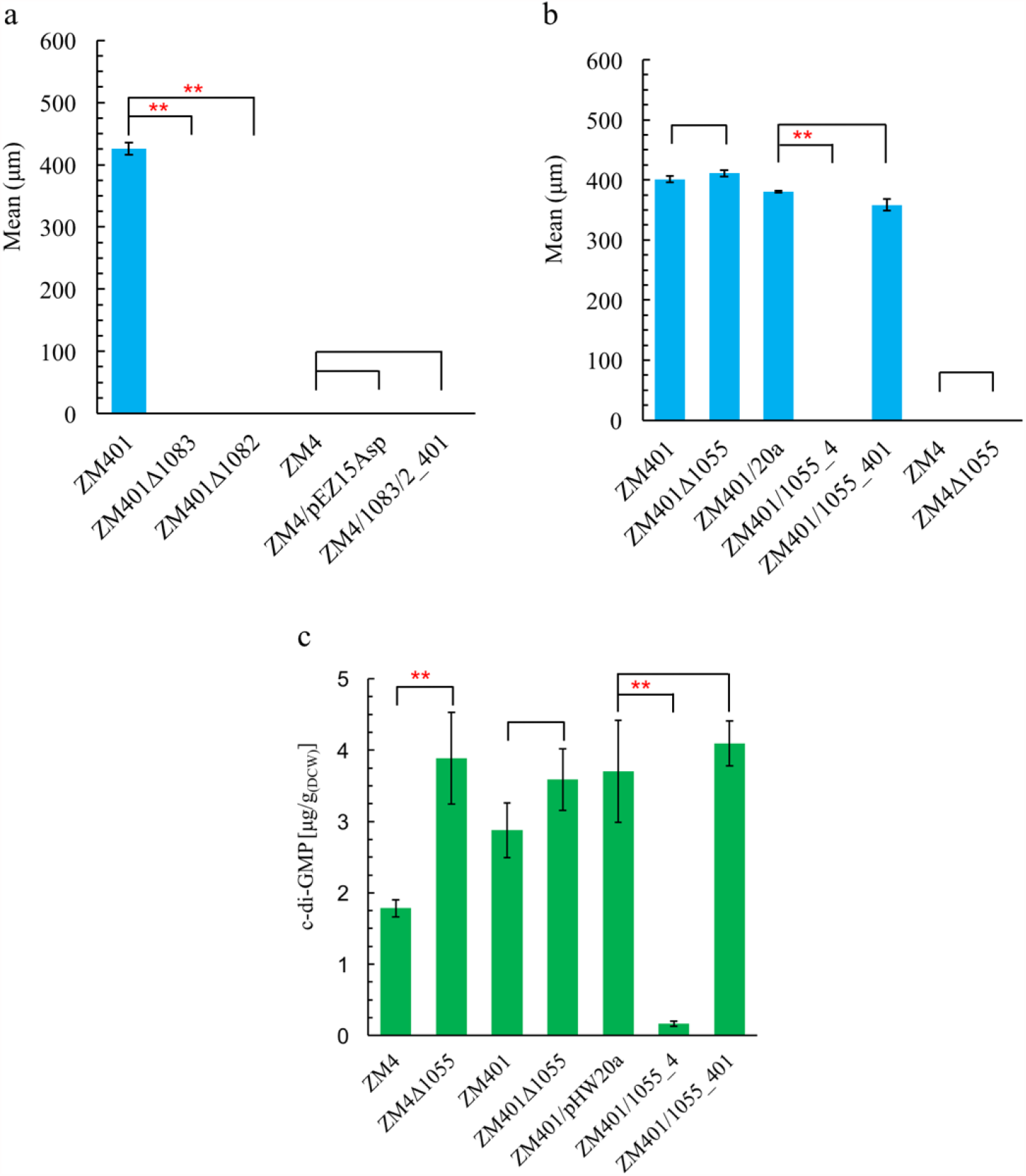
Statistic means for the chord length of ZM401 engineered with the deletion of either ZMO1083 or ZMO1082 and ZM4 engineered with the overexpression of ZMO1083/2 from ZM401 (a). ZM401 and ZM4 engineered with the deletion of ZMO1055, and ZM401 engineered with the overexpression of ZMO1055 from ZM4 and ZM401, respectively (b). Intracellular c-di-GMP analysis for all strains (c). Triplicates were measured, and the significance was analyzed by the *t*-test with *p* < 0.01 (^**^). Raw data for the FBRM analysis is available in the Supplementary Materials: **Figs. S5** and **S6**.

On the other hand, when ZMO1055 from ZM4 was overexpressed in ZM401, its self-flocculation was also disrupted completely with the chord length decreased to the undetectable level, but no significant impact was observed on the self-flocculation of ZM401 when ZMO1055 with the SNP mutation was either deleted or overexpressed (**Fig. 4b**). Meanwhile, no impact on morphology was observed when ZMO1055 was deleted from ZM4. These results indicate that the impact of the SNP mutation in ZMO1055 on the function of the EAL domain encoded by the gene for c-di-GMP degradation is another important factor for the self-flocculation of ZM401.

The intracellular accumulation of c-di-GMP was further measured to validate the impact of the SNP mutation on the function of ZMO1055 (**Fig. 4c**). When ZMO1055 was deleted from ZM4, the intracellular c-di-GMP increased to 3.88 μg/g compared to 1.78 μg/g detected in the wild-type ZM4, which confirmed the activity of the EAL domain encoded by ZMO1055 in the degradation of c-di-GMP. The intracellular c-di-GMP of 2.87 μg/g was detected in ZM401, which is significantly higher than that detected in ZM4, but lower than that detected in ZM4 with ZMO1055 deleted. When ZM401 was engineered with the overexpression of ZMO1055 from ZM4, its intracellular c-di-GMP decreased drastically to 0.17 μg/g, but no significant change in intracellular c-di-GMP was observed when its own ZMO1055 was overexpressed.

These experimental results clearly validated that the SNP mutation in ZMO1055 substantially compromised the function of the EAL domain for c-di-GMP degradation, and the intracellular accumulation of the messenger molecule consequently enhanced the biosynthesis of cellulose microfibrils under the catalysis of the Bcs complex, in particular the BcsA encoded by ZMO1083/2, for the self-flocculation of the bacterial cells from ZM401.

Industrial applications for the self-flocculation of microbial cells such as their self-immobilization within bioreactors for high biomass density to improve productivity and cost-effective biomass recovery by gravity sedimentation depend on their flocculation efficiency, which was characterized experimentally for ZM401 and ZM4 as well as their mutants developed to validate functions of those genes (**Fig. S7**). The chord length measured for the bacterial flocs is well correlated with their flocculation efficiency.

### Engineering unicellular ZM4 with the self-flocculating phenotype

With the roles of ZMO1083/2 and ZMO1055 in the self-flocculation of ZM401 validated experimentally, we engineered unicellular ZM4 through rational design by deleting its ZMO1055 to enhance intracellular accumulation of c-di-GMP, followed by the overexpression of the whole *bcs* operon from ZM401 for the biosynthesis of cellulose microfibrils.

As expected, the engineered strain ZM4Δ1055/BcsABC_401 developed the self-flocculating phenotype, which was characterized by the mean of 568.5 μm for the chord length of the bacterial flocs, and the intracellular accumulation of c-di-GMP up to 2.28 μg/g, higher than that of 1.78 μg/g only detected in the wild-type ZM4, to effectively activate the cellulose synthesis catalyzed by the Bcs complex for the self-flocculation of the bacterial cells (**Fig. 5**). The dynamic gravity sedimentation of ZM4Δ1055/BcsABC_401 is highlighted in Movie 2.

**Fig. 5.**
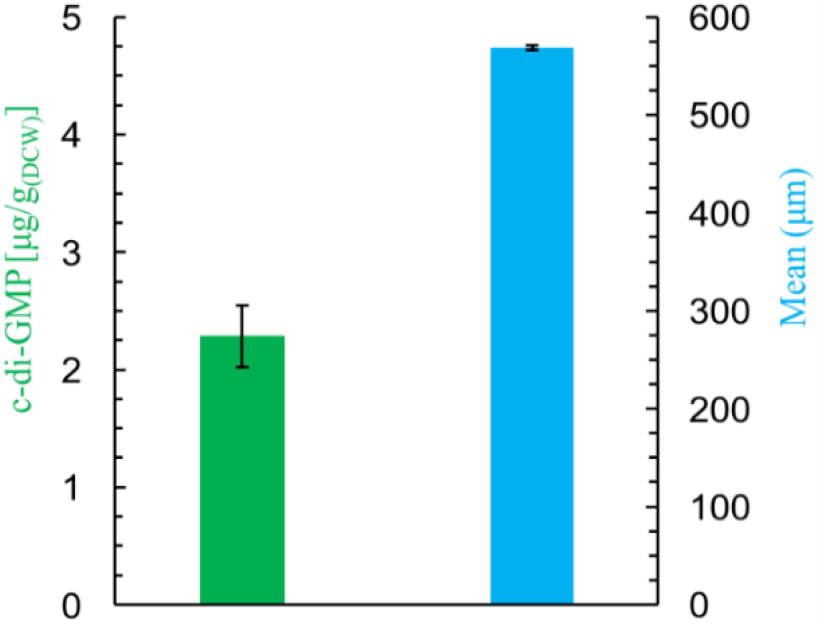
Engineering ZM4 for self-flocculation through the deletion of ZMO1055 and the overexpression of the *bcs* operon from ZM401. The dynamic gravity sedimentation of the bacterial flocs is shown in **Movie 2**, and raw data for their chord length distribution are provided in the Supplementary Materials: **Fig. S6**.

## Discussion

Since ZM4 was sequenced (Seo et al. 2005), many studies have been performed on its genetic modifications to exploit metabolic merits associated with the ED pathway for glycolysis, through which less ATP is produced for lower biomass accumulation, and thus more sugar can be directed to product formation with improved yield (Wang et al. 2018; Liu et al. 2020). Such a trait is very significant, particularly for producing bulk commodities such as cellulosic ethanol and 2, 3-butanediol with major cost from feedstock consumption (Xia et al. 2019; Yang et al. 2016). Unfortunately, less attention has been paid to studies on molecular mechanism underlying the self-flocculation of ZM401, a unique morphology with many advantages for robust production.

Like other bacteria such as *Escherichia coli* (Römling and Galperin 2015), *Z. mobilis* has been predicted to have a *bcs* operon in its genome (Jeon et al. 2012), but almost all strains from this species are unicellular. Some bacteria with *bcs* operons can develop biofilms, but in general surfaces are needed for bacterial cells to attach onto for growth to form stable biofilms (Flemming et al. 2016). Moreover, the matrix of microbial biofilms is composed of extracellular polysubstances, and cellulose could be one of them, but other polymers such as proteins, nucleic acids and lipids are also major components of biofilms (Karygianni et al. 2020). In contrast, the only extracellular polysubstance for the self-flocculation of ZM401 is cellulose, and particularly cellulose microfibrils clearly observed under SEM (Xia et al. 2018).

Bacteria form biofilms with a life cycle: planktonic cells → attachment onto surfaces → biofilm development and maturation → biofilm dispersion (Rumbaugh and Sauer 2020). However a dynamic balance can develop between the self-flocculation of the bacterial cells from ZM401 and the breakup of the bacterial flocs by shearing developed through mixing such as hydrodynamic shearing in shaking flasks and both hydrodynamic and mechanical shearing created by agitation within bioreactors. The larger the bacterial flocs are, the poorer their resistance to shearing will be for being broken up into smaller ones to better resist the shearing. On the other hand, with the growth of the bacterial cells aggregated with the smaller flocs, their size will increase to experience the same fate as those larger flocs. Under suspension culture and fermentation conditions, such a dynamic nature generates size distributions for the bacterial flocs. As a result, their inside can be renewed continuously for sustained viability and metabolic activity, making them more suitable and efficient for industrial production, particularly for continuous culture and fermentation that can be operated for a long time, even yearly without interruption.

Transcriptomic analysis suggested that the *bcs* operon for cellulose synthesis could be responsible for the self-flocculation of ZM401 (Jeon et al. 2012). Not only did experimental studies validate the speculation, but also directly observed and confirmed that cellulose microfibrils make the bacterial cells self-flocculated (Xia et al. 2018), raising such a question: **why the *bcs* operon in ZM4 and all other strains from *Z. mobilis* cannot synthesize cellulose efficiently, and in the meantime make synthesized cellulose developed as microfibrils for the bacterial cells to develop the self-flocculating phenotype and unique morphology with significant advantages for robust production?** Compared to unicellular cells and amorphous biofilms which have been intensively studied, this morphology of bacterial cells has been neglected to a large extent, in particular studies on molecular mechanism underlying the self-flocculating phenotype.

Comparative genomic and transcriptomic analyses and molecular docking simulations for key enzymes and their substrates/activators indicate that the frameshift mutation caused by the nucleotide deletion in ZMO1082 made the putative gene fused with ZMO1083 in ZM401, which substantially altered the structure of BcsA encoded by ZMO1083 in ZM4, particularly its catalytic activity on cellulose biosynthesis. While ZMO1083/2 developed by the fusion of ZMO1082 with ZMO1083 due to the thymine deletion is the necessary condition for synthesizing cellulose microfibrils to flocculate the bacterial cells, another SNP mutation on ZMO1055 is the sufficient condition for such an event to occur, because this mutation compromised the catalytic activity of the phosphodiesterase on c-di-GMP degradation through obstructing the messenger molecule from binding onto the active site of the EAL domain effectively for its intracellular accumulate to active the biosynthesis of cellulose microfibrils.

We therefore develop molecular mechanism for the self-flocculation of *Z. mobile* (**Fig. 6)**. Cellulose is synthesized from UDP-glucose under the catalysis of BcsA encoded by ZMO1083/2, which is activated by the intracellular accumulation of c-di-GMP due to the SNP mutation in ZMO1055, and cellulose microfibrils are further developed and exported by the Bcs complex for the self-flocculation of the bacterial cells.

**Fig. 6.**
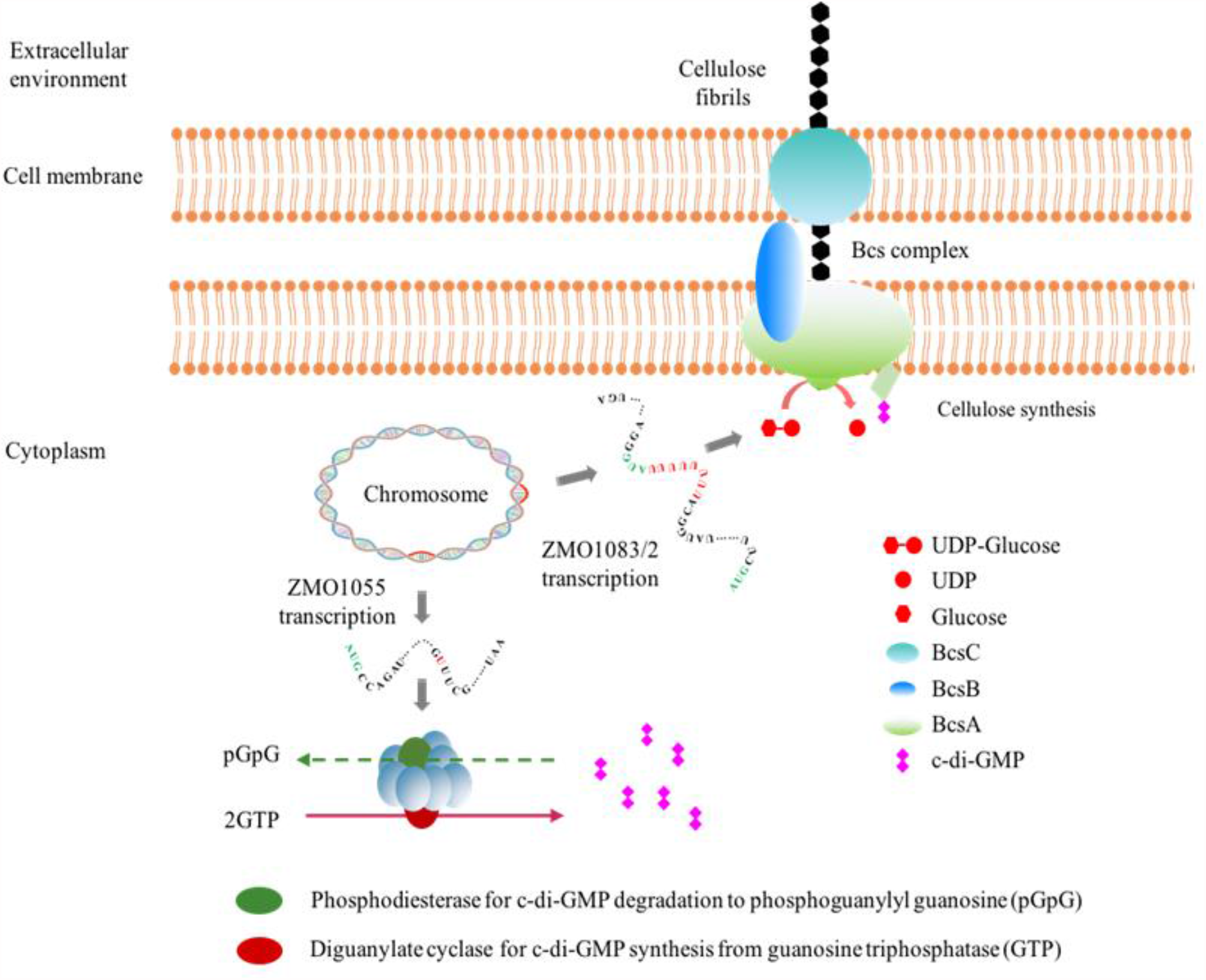
Mechanism underlying the self-flocculation of the bacterial cells from *Z. mobilis*.

Cellulose production from glucose by *Gluconacetobacter xylinum* was discovered more than a century ago (Jacek et al. 2019), and the bioinformatics analyses indicate that ZM4 contains a *bcs* operon comprised of ZMO1083, ZMO1084 and ZMO1085 encoding BcsA, BcsB and BcsC, respectively. We are targeting BcsA encoded by ZMO1083/2 in ZM401 to explore mechanism underlying the biosynthesis of cellulose microfibrils for the self-flocculation of the bacterial cells, since BcsA is the active subunit of bacterial cellulose synthase (Omadjela et al. 2013), and no mutation was detected in other components of the Bcs complex in ZM401.

BcsA in ZM4 contains 665 amino acids, and the topological analysis for the protein indicates that it has 6 transmembrane helices, which seem not enough for cellulose biosynthesis, particularly for synthesizing crystal cellulose as that observed in *G. xylinum* (Jacek et al. 2019). Although the aggregation of ZM4 was reported recently when it was cultured specifically in minimal medium under aerobic conditions, which was attributed to cellulose synthesis (Jones-Burrage et al. 2019), the SEM image shows that the morphology of the bacterial cells is more similar to amorphous biofilms floating in the broth. We therefore reason that BcsA in ZM4 lacks function on catalyzing the synthesis of cellulose effectively, in particularly for synthesized cellulose to develop as microfibrils for the self-flocculation of the bacterial cells.

BcsA in *R. sphaeroides* forms eight transmembrane helices for function to catalyze cellulose biosynthesis (Morgan et al. 2013). With ZMO1082 fused with ZMO1083 in ZM401, the new gene ZMO1083/2 encodes 725 amino acids for two more transmembrane helices, making the BcsA functional not only for cellulose biosynthesis, but also for the development of cellulose microfibrils and their exporting through the coordination of other subunits in the Bcs complex to flocculate the bacterial cells. The number of 725 amino acids for BcsA in ZM401 is close to that of 788 amino acids for BcsA in *R. sphaeroides* (Morgan et al. 2013), and both can catalyze cellulose biosynthesis, but morphologies of synthesized cellulose are significantly different, since *R. sphaeroides* synthesizes amorphous cellulose, raising a necessity for deciphering the structure of BcsA in ZM401 to explore mechanism underlying its catalysis for cellulose biosynthesis, and particularly for the development of cellulose microfibrils to flocculate the bacterial cells.

Another impact on the self-flocculation of ZM401 is from ZMO1055. Although the SNP mutation on ZMO1055 did not change the overall structure of the bifunctional diguanylate cyclase/phosphodiesterase encoded by the gene substantially, the mutation could affect the binding of c-di-GMP onto the EAL domain with the phosphodiesterase, and consequently compromises the degradation of c-di-GMP for more intracellular accumulation of the messenger molecule to activate the cellulose biosynthesis. The intracellular concentration of c-di-GMP in ZM401 engineered with the overexpression or deletion of ZM1055 from ZM401 and ZM4 experimentally supports such a speculation. Therefore, the EAL domain of the protein encoded by ZMO1055 with the SNP mutation in ZM401 directly regulates the cellulose biosynthesis by the Bcs complex, and consequently contributes to the self-flocculation of the bacterial cells.

We therefore propose a more general molecular mechanism for the self-flocculation of bacterial cells: the intracellular biosynthesis of cellulose, development of cellulose microfibrils and their exportation across cell membranes through the expression of *bcs* operon that is functional or heterologous expression of *bcs* operon for the function, and such a process needs to be activated by engineering the bacterial cells with the intracellular accumulation of c-di-GMP, either by deactivating activities of enzymes for its degradation or overexpressing enzymes for its biosynthesis. Under these guidelines, we engineered the unicellular *E. coli* strain MG1655 through overexpressing the gene *wspR* encoding WspR in *Pseudomonas aeruginosa* to catalyze the biosynthesis of c-di-GMP (Liu et al. 2018) as well as the whole *bcs* operon from ZM401 for the biosynthesis of cellulose microfibrils to flocculate the bacterial cells. As expected, experimental results validated our proposed mechanism for the self-flocculation of bacterial cells (**Figs. S8** and **S9**).

Although no significant impact has been observed in growth and ethanol production between ZM401 and ZM4 at present (**Fig. S10**), the self-flocculation of bacterial cells should be controlled properly for industrial applications to avoid potential risk in internal mass transfer limitation on nutrient uptake from medium into flocs and product excretion from the inside into the bulk environment if the self-flocculation of bacterial cells is too strong and large flocs are formed. With the elucidation of molecular mechanism underlying the self-flocculation of ZM401, strategies can be developed for optimizing the self-flocculating process. Moreover, theoretically and technically, other bacteria can also be engineered with the phenotype for robust production.

## Methods

### Strains and media

Strains used in this work are presented in **Table S3**. Rich media RMG2 (10 g/L yeast extract, 2 g/L KH_2_PO_4_ and 20 g/L glucose) and RMG10 (10 g/L yeast extract, 2 g/L KH_2_PO_4_ and 100 g/L glucose) were used for seed culture and ethanol fermentation with *Z. mobilis*, respectively. Luria-Bertani (LB) medium was used for the culture of all *E. coli* strains, such as *E. coli* DH5α and *E. coli* JM110. All media were sterilized by autoclaving at 115 °C for 20 min before inoculation.

### *Z. mobilis* culture and ethanol fermentation

A single colony of ZM401 on agar plate was inoculated into 5 mL RMG2, which was cultured at 30 °C and 150 rpm for 24 h, and then used for DNA extraction and genome sequencing.

A loopful of ZM401 or ZM4 on agar plate was inoculated into 150 mL Erlenmeyer flask containing 100 mL RMG2 for culture at 30 °C and 150 rpm till OD_600_ increased to ∼ 1.0 as seed culture. The fermenter (KoBioTech KF-2.5L, Korea) containing 1.4 L RMG10 was inoculated with 100 mL seed culture for a total working volume of 1.5 L, and ethanol fermentation was performed at 30 °C and 150 rpm with pH controlled at 6.0 by automatic titration by 2 N KOH for comparison with the transcription analysis previously reported for ZM4 (Yang et al. 2009b).

We experimentally observed that microaeration benefits the self-flocculation of ZM401 without significant impact on its growth and ethanol production. Therefore, air filtered with 0.2 μm membrane was sparged into the fermenter at an aeration rate of 0.1 L/min, equivalent to 0.067 vvm, volumetric aeration rate per unit working volume of bioreactors and per minute, to stimulate the self-flocculation of ZM401 for mining more information on its gene expression associated with the self-flocculating phenotype of the bacterial cells.

### Analysis for cell growth and ethanol fermentation

The culture of 4 mL was sampled and de-flocculated by cellulases at room temperature for complete de-flocculation following the protocol developed previously with minor modifications (Xia et al. 2018), which was then vortexed vigorously for homogenous suspension, followed by OD_600_ measurement using the Microplate Reader (Multiskan GO, Thermo Fisher, USA) to characterize cell density. In case of need, the cell density characterized by OD_600_ was converted to dry cell weight (DCW).

Glucose, ethanol and glycerol were analyzed by HPLC (Waters e2695, USA) with the RI detector (Waters 2414, USA) operated at 50 °C. An organic acid column (Aminex HPX-87H 300 × 7.8 mm, Bio-Rad, USA) was employed for the analysis, which was operated at 65 °C with 10 mmol/L H_2_SO_4_ as the flow phase at a flow rate of 0.6 mL/min.

### In situ characterization for the self-flocculation of *Z. mobilis*

Unlike unicellular cells, bacterial flocs are deformable and not uniform in size, which cannot be characterized directly through microscope observation, and online measurement is needed for characterizing their size distributions.

The floc chord length counts per second were detected by FBRM (ParticleTrack® G400, Mettler Toledo, USA) and analyzed by iC FBRM 4.4 as previous study with minor modifications (Ge et al. 2005). The bacterial flocs cultured in shaking flasks near the end of their exponential growth phase were collected, and transferred into the container equipped with a stirrer for analysis using FBRM. The FBRM sensor was scanned at a speed of 2 m/s with an interval of 10 s. The stirrer was adjusted to a rotating rate of 150 rpm for homogenous suspension of the bacterial flocs. The FBRM analysis was run for at least 15 min with each sample to make the data set statistically reliable.

The chord length of a single floc was calculated automatically through multiplying the time difference detected for the laser beam to be reflected by the bacterial floc by the scanning speed. Data were analyzed and processed using Macro V. 1.1.11 with smoothness for every 5 measurements. Square weight mean was used to assess the statistical data. Counts less than 5 were marked as undetectable.

### Measurement of flocculation efficiency for the bacterial cells

Flocculation efficiency was measured and calculated as previously reported with some modifications (Xia et al. 2018). OD_600_ measurement for cell density as described above was designated as A. The culture collected simultaneously as that for the cell density analysis was rested statically for 5 min, and then 400 μL supernatant was sampled, which was treated with cellulases to de-flocculate any small flocs suspended in the supernatant, and vortexed for homogenous suspension to measure OD_600_ as B. The flocculation efficiency (F) was calculated by the equation:

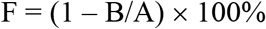

### Genomic DNA extraction and genome sequencing

The genomic DNA was extracted from 5 mL culture inoculated with the single colony of ZM401 using the Bacterial DNA Kit (OMEGA, USA).

The genome sequencing was performed using the 3^rd^ generation sequencing PacBio RS II System at Instrumental Analysis Center, Shanghai Jiao Tong University (Shanghai, China). The Next Generation Sequencing (NGS) was performed using Illumina X Ten platform at GENEWIZ, Inc. (Suzhou, China) by the paired-end sequencing technology according to the standard protocols.

As a mutant from ZM4, the SNVs of ZM401 were identified with the genome of ZM4 as the reference (GenBank accession numbers: NZ_CP023715 for the chromosome, and NZ_CP023716, NZ_CP023717, NZ_CP023718 and NZ_CP023719 for the plasmids), which combined the results from the CLC Genomics Workbench (version 11.0) with NGS data and the bcftools’ (Version 1.9) SNPs calling function with the input of PacBio data. The structure variations (SVs) were analyzed with the long-reads PacBio data based on previously reported methods (Sedlazeck et al. 2018). Briefly, NGMLR (https://github.com/philres/ngmlr) was used for aligning the long-reads, and Sniffles (https://github.com/fritzsedlazeck/Sniffles) was used for the SV identification.

### Transcriptomic analysis

The transcriptomic analysis workflow was referred to previous work (Yang et al. 2009b). Briefly, cultures were sampled from the fermenter at 6 h when the bacterial cells entered into their exponential growth stage, followed by total RNA extraction using the TRIzol reagent (Invitrogen, USA). rRNA was depleted using the Ribo-off rRNA Depletion Kit (Bacteria NR407). The sequencing library was then constructed after connecting the sequencing adapters with the cDNA fragments, of which the first-strand cDNA was synthesized using random hexamer-primers and the second-strand cDNA was synthesized using dATPs, dGTPs, dCTPs, dUTPs, RNase H and DNA polymerase I respectively after removing dNTPs.

RNA-Seq was performed using the paired-end sequencing technology according to the standard Illumina protocol at GENEWIZ, Inc (Suzhou, China). The quality of the RNA-Seq fastq data was evaluated using the FastQC program (www.bioinformatics.babraham.ac.uk/projects/fastqc/). Data passing the quality control were imported into the CLC Genomics Workbench (version 11.0, Qiagen, USA) for the RNA-Seq analysis to obtain the RPKM (reads mapping to the genome per kilobase of transcript per million reads sequenced) values using the genome of ZM4 as the reference. Gene expression normalization, analysis of variance (ANOVA) and hierarchical clustering analysis were conducted using JMP Genomics (version 9.0, SAS Inc, USA) to identify differentially expressed genes in ZM401 compared to ZM4 with the selection thresholds −1 ≥ Log_2_(Fold Change) ≥ +1 and the statistical significance *P* ≤ 0.05. Duplicate samples were used for each analysis.

### Protein-ligand docking

Protein structures were constructed by SWISS-MODEL (Waterhouse et al. 2018) with BcsA from *R. sphaeroides* (Morgan et al. 2013) and RbdA from *P. aeruginosa* as the references (Liu et al. 2018), which were pre-processed by Prepare Protein in Discovery Studio (2016, BIOVIA). Then substrates, products and activators were pre-processed by Prepare Ligands, and docked with the proteins in Discovery Studio (2016, BIOVIA). Interactions between proteins and those molecules were simulated and analyzed through protein-ligand docking.

### Genetic manipulations

Plasmids and primers used in this study are listed in **Table S4** and **Table S5**, respectively. Plasmid construction and genetic manipulations were performed as previously reported (Xia et al. 2018). For gene deletion, two homologous regions flanked target gene were obtained by PCR, then infused with the plasmid pEX18Tc. For gene overexpression, target genes and promoters were amplified by PCR, and then infused with the plasmid pHW20a or pEZ15Asp. Recombinant plasmids were transformed into *E. coli* DH5α for propagation, which were confirmed by Sanger sequencing before being transformed into *E. coli* JM110 for demethylation. Demethylated recombined plasmids were then transformed into ZM401 or ZM4 through electroporation using Gene Pulser (Bio-Rad, USA) with 0.1 cm gap cuvettes operated at 18 kV/cm, 200 Ω and 25 μF. Selection of positive colonies was carried out on the RMG2 agar plate containing 20 mg/L tetracycline or 100 mg/L spectinomycin for mutants engineered with the vector pHW20a or vector pEZ15Asp. While engineered with the vector pEX18Tc, colonies were screened on RMG2 agar plate supplemented with 20 mg/L tetracycline followed by another selection on RM agar plate (rich medium composed of 10 g/L yeast extract and 2 g/L KH_2_PO_4_ but without glucose) supplemented with 50 g/L sucrose.

### Extraction and measurement of intracellular c-di-GMP by LC-MS

The bacterial cells were grown to their exponential phase with OD_600_ ∼3.0, and then 15 mL culture suspension was centrifuged at 5 000 g and 4 °C for 3 min to collect the pellet. The pre-cooled extraction solvent (2:2:1 for methanol: acetonitrile: ddH_2_O) of 1 mL was added to disperse the pellet. The suspension was maintained at −20 °C for 30 min to disrupt the bacterial cells, and also extract soluble components including c-di-GMP, which was then centrifugalized at 12 000 g and 4 °C for 5 min to collect the supernatant. The pellet was dispersed again with 1 mL pre-cooled extraction solvent for another round of extraction, and the supernatant was combined with that collected from the first round of extraction for further treatment.

Chloroform of 2 mL was added into the supernatant (∼2 mL), which was mixed through hand-shaking to extract hydrophobic components to remove impurities. After resting 5 min for phase separation, the upper phase with c-di-GMP was collected for drying by a freeze dryer. The extraction solvent of 80 μL was used to dissolve the paste, which was vortexed violently for 2 min, and then 50 μL solution was sampled for c-di-GMP analysis.

c-di-GMP separation and quantification were conducted by Acquity UPLC I-class/VION IMS QTOF. Briefly, Waters I-Class Acquity UPLC (Waters, UK) with SeQuant ZIC-HILIC column (100 mm × 2.1 mm) packed with 3.5 μm polyetheretherketone (Merck, Germany) was used for separating c-di-GMP, which was operated at 45 °C. The flow phase was developed by Phase A: 50 mM ammonium formate and Phase B: acetonitrile. Separation of the samples was conducted at three stages within 15 min at a flow rate of 0.4 mL/min: Stage I from 0 to 10 min with Phase B decreased from 90% to 50% and Phase A increased from 10% to 50%, Stage II from 10 min to 12 min with Phase B increased from 50% to 90% and Phase A decreased from 50% to 10%, and Stage III from 12 min to 15 min with 90% Phase B + 10% Phase A. Separated substances were further treated by Vion IMS QToF mass spectrometer (Waters, UK) with an electrospray ionization (ESI) interface operated at negative ion mode with parameters: desolvation temperature, 500 °C; ion source temperature, 120 °C; collision energy, 40 eV; desolvation gas (nitrogen) flow rate, 1000 L/h; cone gas flow rate, 50 L/h; capillary voltage, 2000 V; cone voltage, 20 V. Data were acquired from m/z 50-1000 at a scan speed of 0.3 s with the scan mode mRm. UNIFI 1.8.1 was applied for data processing. c-di-GMP with a purity of 98% (Biolog, Germany) was used as the standard to calibrate the analysis.

### Engineering *E. coli* K12 with the self-flocculation phenotype

A commonly used *E. coli* K12 strain (MG1655) was selected for engineering with the self-flocculating phenotype through rational design developed based on the understanding of mechanism underlying the self-flocculation of *Z. mobilis*.

Plasmids and primers for engineering MG1655 are listed in **Table S5** and **Table S6**, respectively. The plasmid pA2c was engineered previously with the *bcs* operon from ZM401 for expression in MG1655 (Jia et al. 2020). The gene *wspR* cloned from *Pseudomonas aeruginosa* and the plasmid pE1k were amplified by PCR, and then ligated using the Golden Gate cloning (Engler et al. 2008) for amplification in *E. coli* DH5α to transform MG1655 and enhance its biosynthesis of c-di-GMP. The transformation of MG1655 with the pE1k and both the pA2c and pE1k was performed by electroporation using Gene Pulser (Bio-Rad, USA) with 0.1 cm gap cuvettes operated at 18 kV/cm, 200 Ω and 25 μF, and the transformants were selected on the LB medium containing 50 mg/L kanamycin and 30 mg/L chloramphenicol + 50 mg/L kanamycin, respectively, followed by PCR amplification and DNA sequencing for verification. Colony of MG1655 engineered with pE1k-wspR or pA2c-bcs + pE1k-wspR was inoculated into tube containing 5 mL LB medium for overnight culture at 37 °C and 200 rpm. Then, the culture suspension of 1 mL was transferred into flask containing 100 mL LB medium supplemented with 2 g/L glucose, which was incubated at 37 °C and 200 rpm to the mid-log phase with OD at 0.6∼0.8. The expression of *wspR* and *wspR* + *bcs* was induced by supplementing isopropyl β-D-1-thiogalactopyranoside (IPTG) to 0.1 mM and both IPTG and anhydrotetracycline to 0.2 mg/L, and the induction was performed at 25 °C and 150 rpm, which was lasted for 4 days.

### Data access

All data are available in the main text and the Supplementary Materials. Raw data for omics analysis were deposited at National Center for Biotechnology Information (NCBI) with the BioProject accession numbers of PRJNA590867 and PRJNA590816 for genome sequencing/annotation and transcriptomic analysis, respectively. Upon request, all materials developed in this work are available for non-commercial purpose.

## Acknowledgements

We are grateful to Prof. Xiaoxia Xia for donating the model strain *Escherichia coli* K12 (MG1655) and Dr. Ying Xu, Prof. Yi Xiao and Prof. Jie Bao for donating the plasmids. This work was financially supported by National Natural Science Foundation of China with the grant numbers of 21536006 and 31970026.

